# Lineage-specific patterns of sexually dimorphic antennal transcription in the paper wasp *Polistes fuscatus*

**DOI:** 10.1101/2024.07.30.605902

**Authors:** Andrew W. Legan, Michael J. Sheehan

## Abstract

Olfaction mediates many behaviors in social Hymenoptera, with sexual dimorphism in antennal transcription associated with different behaviors between sexes. Females display coordinated social behaviors within colonies, while males exhibit limited social behavior but are selected for finding mates. The expanded “9-exon” odorant receptor (OR) gene subfamily is associated with chemical communication and exhibits strongly female biased antennal transcription in ants and honey bees. Polistine wasps represent an independent evolution of sociality and associated expansion of 9-exon ORs, though antennal expression patterns are unknown. Here, we report distinct patterns of sexually dimorphic OR transcription in *Polistes fuscatus* compared to ants and bees. Most *P. fuscatus* 9-exon transcripts were detected at similar levels in males and females, and some were male biased. We also report differential antennal transcription of cytochromes P450 and muscle-related genes between sexes. We discuss these patterns in the context of the unique sexual and social behaviors of *Polistes* wasps, including prolonged male mating aggregations and male antennal tapping and curling during courtship and copulation. These results call attention to the lineage-specific selective pressures shaping sexually dimorphic antennal transcription in social insects.

## Introduction

Selective pressures frequently differ between sexes, leading to sex-specific fitness peaks shaping behaviors and sensory abilities across diverse animal lineages (Lande 1980; Parsch and Ellegren 2013). Among insects, selection favoring improved abilities to search for mates has led to substantial investment in sensory organ size and acuity in the courting sex (Rospars and Chambille 1989; Nakanishi et al. 2009; Zhao et al. 2016; Scherberich et al. 2017). Many male insects with visual mate search behavior have larger eyes with larger facet size compared to females (Beersma et al. 1977; Gullefors and Petersson 1993; Ziemba and Rutowski 2000; Somanathan et al. 2017). In insect species in which female sex pheromones attract male mates, more olfactory sensilla cover male antennae than female antennae (Chapman 1982; Hansson and Stensmyr 2011). In the honey bee *Apis mellifera*, drone (male) antennae are longer and more densely populated with queen pheromone-sensitive olfactory sensilla compared to female antennae, aiding in olfactory localization of queens during mating flights of competing drones (Brockmann and Brückner 2001; Koeniger et al. 2005; Wanner et al. 2007b). Sexual dimorphism is extensively documented in moth species, with male to female antennal surface area ratios of 5:1 in *Antheraea pernyi*, and 2:1 in *Manduca sexta* (Kaissling 1971; Sanes and Hildebrand 1976; Lee and Strausfeld 1990).

In addition to peripheral sensory morphology, chemoreceptor proteins contribute to variation in olfactory behavior between individuals (Rihani and Sachse 2022). Sexual selection may manifest at the molecular level in sexually dimorphic transcription of genes encoding chemoreceptor proteins (Zhou et al. 2009; Andersson et al. 2014; Zhang and Löfstedt 2015; Bastin-Héline et al. 2019). For example, in the silk moth *Bombyx mori*, host plant-seeking by females is mediated by the female biased odorant receptors *BmOr19* and *BmOr30*, and mate-seeking by males is mediated by the male biased *BmOr1* and *BmOr3* (Wanner et al. 2007a; Nakagawa et al. 2005). In sum, chemoreceptors are essential for sex-specific olfactory behavior in diverse insect lineages. However, sexually dimorphic odorant receptor transcription, or antennal gene transcription more generally, has not been previously investigated in social wasps. To fill this gap, we studied sexual dimorphism of antennal gene transcription in the paper wasp *Polistes fuscatus* and compared our results with patterns observed in ants and social bees.

In social Hymenoptera, contrasts in sexually dimorphic selective pressures are especially stark. Colonies function by the collective behavior of females, and males usually contribute little to the social coordination of the colony (Boomsma et al. 2005). Instead, males typically eclose late in the colony cycle and are short-lived, focusing almost exclusively on reproduction. For example, in *A. mellifera*, the hallmark of male behavior is its “simplicity” and “limited diversity” (Ohtani 1974). In light of their life history differences, it is not surprising that sex differences in sensory morphology between male and female social insects have been widely reported (Brockmann and Brückner 2001; Renthal et al. 2003; Koeniger et al. 2005; Wanner et al. 2007b; Heinze et al. 2021).

Expanded sets of odorant receptor (OR) genes have evolved repeatedly among social Hymenoptera, giving rise to perception of complex chemical cues and signals (Robertson and Wanner 2006; Smith CR, Smith CD et al. 2011; Smith CD, Zimin A et al. 2011; Zhou et al. 2012; Engsontia et al. 2015; Zhou et al. 2015; Karpe et al. 2016; McKenzie et al. 2016; Legan et al. 2021). Hymenopteran odorant receptors are divided into subfamilies based on their sequence similarity (Zhou et al. 2012). In the honey bee and some ant species, these subfamilies are associated with putative functions based on electrophysiological responses of the encoded ORs to chemical stimuli (Wanner et al. 2007b; Pask et al. 2017; Slone et al. 2017). The “9-exon” subfamily is of particular note because these receptors respond to important social chemical cues, cuticular hydrocarbons (CHCs) (Pask et al. 2017; Slone et al. 2017). Given observations of limited social behavior in male social Hymenoptera, males would be predicted to express a limited range of odorant receptors attuned to social chemicals like CHCs. Indeed, antennal gene transcription of 9-exon subfamily ORs is strongly upregulated in female workers relative to males in ants and bees (Zhou et al. 2012; McKenzie et al. 2016; Karpe et al. 2016). To the contrary, an odorant receptor in *A. mellifera* (*AmOr11*) which responds to components of the queen mandibular pheromone exhibits male biased transcription (Wanner et al. 2007b). *AmOr11* is in the “L” subfamily of Hymenopteran odorant receptors. In *A. mellifera* and *Apis florea*, antennal transcription of L subfamily ORs is biased towards drones compared to workers (Wanner et al. 2007b; Karpe et al. 2016). It is unknown whether independent evolution of sociality in wasps has coincided with similar patterns of sexual dimorphism in OR subfamily transcription.

Features of the *Polistes* mating system and male behavior make paper wasps an intriguing clade for the study of sexual selection (Beani et al. 2014). Nest founders and workers are all female. Reproductive females (gynes) emerge in the late summer and early fall, mate, and overwinter before founding nests in the spring (West-Eberhard 1969; Reeve 1991; Miller et al. 2018). These gynes do not provision brood and are typically found on the nest, where they invest less in nest defense behavior than workers (West-Eberhard 1969; Reeve 1991; Judd 2000). Males emerge concurrent with gynes (West-Eberhard 1969). In the context of the nest, male *Polistes fuscatus* and *Polistes metricus* will sometimes feed larvae, a behavior typically assumed to be expressed only in workers and queens, but they have not been observed to hunt for caterpillar prey to feed larvae (Hunt and Noonan 1979). *Polistes* males employ multiple mate search strategies, including aggregation, territory defense, and ranging (Beani and Turillazzi 1988; Beani et al. 2019). In multiple *Polistes* species, males aggregate in groups, where they await mating opportunities from visiting females (Post and Jeanne 1983a, 1983b, 1984; Beani et al. 1992; Polak 1992; de Souza et al. 2017b; Mov. S1A; Fig. 1A). Males can spend weeks in such groups, providing an unusually prolonged and rich social life compared to males in other social insect species (Fig. 1A). Overall, the different social environments occupied by female and male *Polistes* wasps may result in sexually dimorphic selective pressures shaping their chemosensory system.

**Figure 1.**
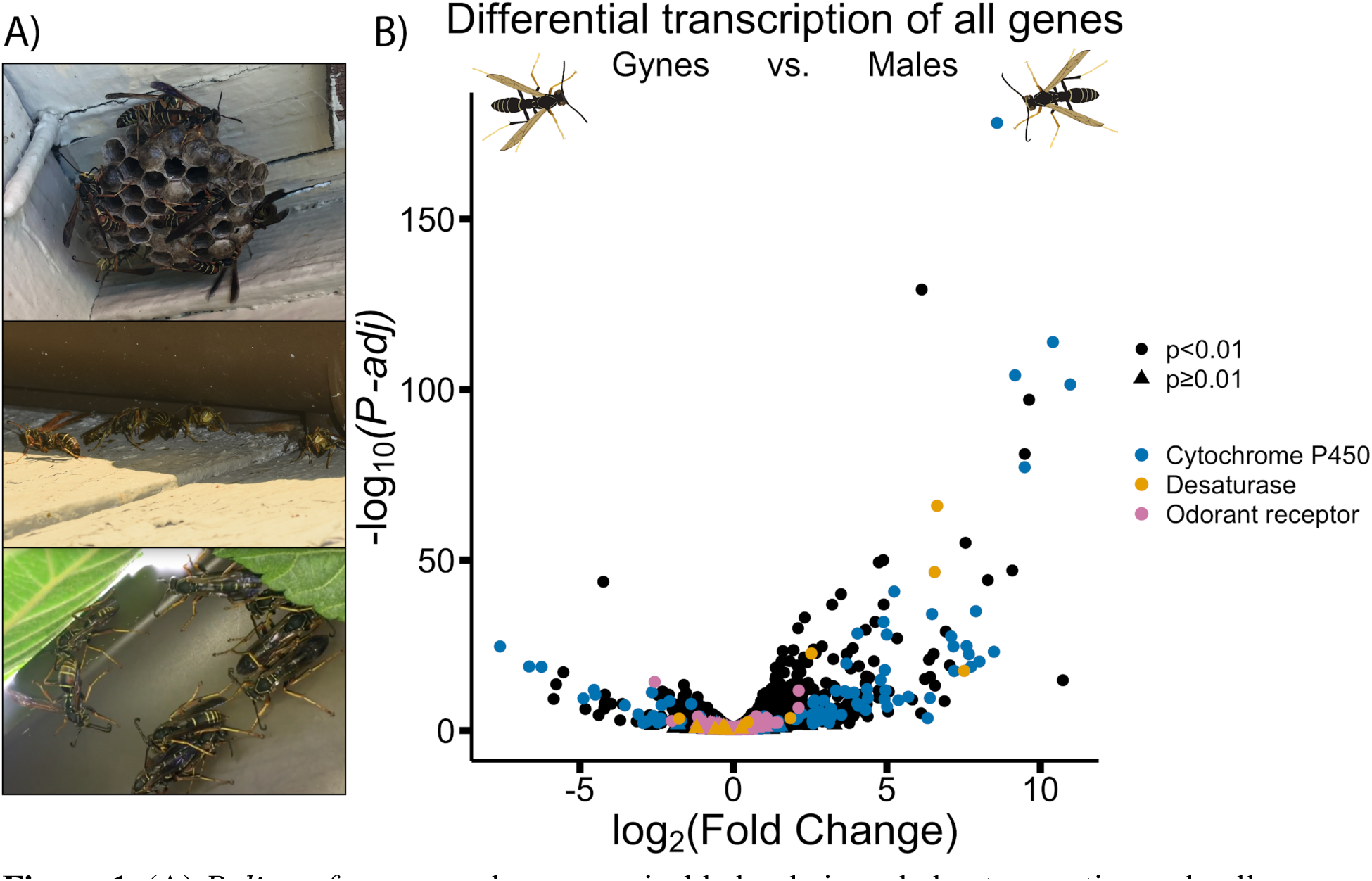
(A) *Polistes fuscatus* males, recognizable by their curled antennae tips and yellow faces, eclose in the late summer and early fall and can be observed on their natal nest (top) as well as in groups of males aggregated away from the nest (middle, bottom). (B) Volcano plot showing sexually dimorphic antennal gene transcription of all detected genes. Positive fold change values were biased towards male antennae, and those with negative fold change values were biased towards gyne antennae. Genes represented by circles were differentially transcribed at Benjamini and Hochberg adjusted significance threshold of *P-adj* < 0.01.

There is evidence that *Polistes* gynes and males utilize olfaction to find and assess potential mates. In *Polistes exclamans*, both males and gynes are attracted to chemicals of the opposite sex (Reed and Landolt 1990; Elmquist et al. 2018). Male *Polistes dominula* likely discriminate between workers and gynes, and between parasitized and non-parasitized gynes, based on chemical cues (Cappa et al. 2013). These results suggest that in *Polistes*, both males and females rely on olfaction to find and assess mates. However, the mate attractant chemicals may differ between sexes, as there is moderate sexual dimorphism in cuticular hydrocarbons between male and female paper wasps (Espelie and Hermann 1990; Layton et al. 1994; Beani et al. 2019; de Souza et al. 2022). There is also some sexual dimorphism in the shape and composition of *Polistes* antennae. In *Polistes dominula*, male antennae house several secretory glands absent in female antennae (Romani et al. 2005). Male *Polistes* have an additional antenna segment compared to females and are typically equal or smaller in body size compared to females (Miller and Sheehan 2021). Furthermore, distal segments of *Polistes* male antennae curl to form a hook at the antenna terminus (Pekkarinen and Gustafsson 1999). During courtship, males tap females with their antennae and curl their antennae around the female’s antennae during copulation (West-Eberhard 1969; Romani et al. 2005; Mov. S1B).

Given the genus-wide sexual dimorphism of antennal morphology, antennal gland composition, CHC profiles, mate search behavior, and courtship antennation, selection has likely favored sex-specific olfactory perception in *Polistes*. *Polistes* genomes encode ∼200 putative odorant receptor genes, and the 9-exon OR subfamily makes up about half of all candidate *Polistes* ORs (Legan et al. 2021). The L subfamily is also a considerable size at 24 genes, all within an ancient tandem array ancestral to ants, bees, and wasps (McKenzie and Kronauer 2018; Legan et al. 2021). Considering the patterns of sexually dimorphic transcription of 9-exon and L subfamily ORs in ants and honey bees, we anticipated similar patterns in social wasps as a result of evolutionary convergence during social evolution. To test this prediction, we sequenced the antennal mRNA of the northern paper wasp *P. fuscatus*. We compared males to gynes, eliminating the confound of reproductive status present in comparisons between males and workers. We report general patterns of sexual dimorphism in antennal gene transcription before focusing specifically on the odorant receptors.

## Materials and Methods

### RNA sequencing of *P. fuscatus* antennae

Six *P. fuscatus* males and six reproductive females (gynes) were collected in the fall of 2017 in Ithaca, NY. Each wasp was collected from a different nest, except one female and one male that were collected from the same nest. Wasps were housed in deli cups in the lab, with water vials and sugar cubes provided *ad libitum*, and were provided an opportunity to mate.

Antennae were resected from heads and flash frozen in liquid nitrogen, then stored at -80°C until RNA isolation using the Qiagen RNeasy Mini kit according to the manufacturer’s recommendations (QIAGEN Sciences, Germantown, MD, USA). Each RNA sample represented the total RNA isolated from two pooled antennae of each individual. Libraries were prepared in house using the NEBNext Ultra II RNA Library Prep Kit for Illumina #E7770L with the NEBNext Poly(A) mRNA Magnetic Isolation Module #E7490 and NEBNext Multiplex Oligos for Illumina #E7600S (New England Biolabs, Ipswich, MA, USA) and sequenced on two lanes of an Illumina HiSeq at Novogene (Davis, CA, USA). Raw reads were trimmed using Trimmomatic v.0.39 (Bolger et al. 2014) as described in Legan et al. (2021). Trimmed reads (average length 132 bp) were then mapped to the *P. fuscatus* genome (NCBI RefSeq GCF_010416935.1; Miller et al. 2020) using STAR v.2.7.9 (Dobin et al. 2013).

### Statistical analyses of antennal transcription in *P. fuscatus*

The bedtools multicov() function was used to generate read counts per gene, treating all reads as single end reads and counting reads mapping to exons (Quinlan and Hall 2010). Sexually dimorphic transcription was quantified in R version 3.5 and R version 4.2.2 using the edgeR package (version 3.40.0) (Robinson et al. 2010; R Core Team 2018). Because male antennal RNA transcription sequenced in sample “M50” (SAMN34203595) was an extreme outlier in a non-metric multidimensional scaling (NMDS) analysis (Fig. S1), this sample was excluded from analysis. The edgeR glmLRT() function was used to test for differential transcription of genes using likelihood ratio tests (Chen et al. 2008). *P*-values reported represent Benjamini and Hochberg adjusted *P*-values (Benjamini and Hochberg 1995), output of the edgeR function topTags(). We report these values as *P-adj* in the text. Gene ontology (GO) analysis was done using the topGO package in R using the “classic” method and Fisher’s exact test (Alexa and Rahnenfuhrer 2021). Statistical comparisons of transcription levels between *P. fuscatus* OR subfamilies were conducted using Wilcoxon rank sum tests (Mann-Whitney tests) with the wilcox.test() function in R.

### Phylogenetic reconstruction of *P. fuscatus* odorant receptors

Odorant receptor gene and pseudogene models encoding proteins with lengths less than 350 amino acids were excluded from phylogenetic reconstruction. Protein sequences were aligned using MAFFT (Katoh and Standley 2013) version 7.453 with options “--genafpair” and “--maxiterate 1000” (strategy E-INS-i). The alignment was trimmed using trimAl with option “-gappyout” to remove gap rich columns, generating a phylip alignment of 203 sequences with length 334 amino acids (Capella-Gutiérrez et al. 2009). A phylogenetic tree of odorant receptor protein sequences was constructed with RAxML version 8.2.12 using the maximum likelihood optimality criterion with 100 bootstrap inferences and the odorant receptor coreceptor as outgroup (Jones et al. 1992; Stamatakis 2014). The tree was visualized in R using the ggtree package and transcription levels mapped with the ggtree gheatmap() function (Yu et al. 2017).

### Comparisons between Hymenoptera species

Antennal RNAseq data describing sexually dimorphic odorant receptor transcription in four other Hymenoptera species were accessed in online supplementary materials of the associated manuscripts. Data from the ants *Camponotus floridanus*, *Harpegnathos saltator*, and *Ooceraea biroi* were accessed from supplementary table S7 in Zhou et al. (2015). Data from the honey bee *Apis florea* were accessed from supplementary table Suppl-2 in Karpe et al. (2016). For the ants and bee data, the sign of fold change values was modified so that all datasets had increased male transcription as the positive direction. Phylogenetic reconstruction of Hymenopteran ORs in 9-exon and L subfamilies was carried out using the same parameters for alignment, trimming, and maximum likelihood tree creation as described above, with no outgroup specified. Amino acid sequences for the honey bee *Apis florea* were accessed from supplementary file Suppl-1 in Karpe et al. (2016) and protein sequence data for two ants, *C. floridanus* and *H. saltator*, were accessed from Dataset S1 in Zhou et al. (2012). Only those odorant receptors with predicted protein sequences greater than 350 amino acids in length were included in phylogenetic reconstruction. Statistical comparisons of subfamily-specific proportions of sexually dimorphic ORs between Hymenoptera species were conducted using Fisher’s exact tests with the fisher.test() function in R. Statistical comparisons of sexually dimorphic transcription of 9-exon and L subfamily ORs (as represented by log_2_(fold change) values) between *P. fuscatus* and the four other species were conducted using Wilcoxon rank sum tests (Mann-Whitney tests) with the wilcox.test() function in R.

## Results

### Sexually dimorphic antennal transcription of CYPs, muscle-related, and other genes in *P. fuscatus*

*P. fuscatus* adults were distinctly separated by sex along the first MDS axis, which explained 44% of the variation in antennal gene transcription between samples (Fig. S2). Of 12,155 predicted gene models, 77% (9,395 genes) were detected at CPM greater than one in at least four samples (hereafter “transcribed genes”). A subset of 1,077 genes (11.5% of transcribed genes) were significantly differentially transcribed between sexes (*P-adj* < 0.01; Fig. 1B; Fig. S3). In both males and females, the top differentially transcribed gene was a cytochrome P450 (CYP). Of the 205 predicted CYP gene models in the *P. fuscatus* genome, 93 were significantly differentially transcribed at *P-adj* < 0.01. Of the top 50 male- or female-biased genes, 26 (52% of male biased) and 15 (30% of female biased) were predicted CYPs (Fig. 1B). Three of the top 50 male biased genes were fatty acid desaturases. Of 14 fatty acid desaturase gene models in the *P. fuscatus* genome, eight were differentially transcribed at *P-adj* < 0.01, and six of these were male biased (Fig. 1B). Gene ontology (GO) term analysis identified 17 terms (of 3,031 terms annotated) significantly enriched for 317 differentially transcribed genes at *P* < 0.01, including six terms (89 genes) associated with muscle tissue (Table S1). Long-chain fatty acid metabolic process genes were also enriched in the GO analysis. Examination of these six genes revealed three proteins associated with elongation of very long chain fatty acids, two 4-coumarate-CoA ligase 1-like genes, and one long chain fatty acid CoA ligase ACSBG2.

### Patterns of gene transcription and sexual dimorphism in the *P. fuscatus* odorant receptor gene family

The bulk of the odorant receptor genes displayed average transcription levels in comparison with the rest of the annotated genes in the *P. fuscatus* genome (Fig. S3). This is not surprising given that each OR is expressed in only a subset of the odorant receptor neurons, which in aggregate is only a fraction of the total cells in the antennae. The odorant receptor coreceptor was the most transcribed gene in the odorant receptor gene family (mean CPM = 814 ± 145 SD), consistent with its function as a requisite receptor subunit expressed ubiquitously in olfactory receptor neurons. The top transcribed tuning odorant receptor was in the J subfamily, *PfusOr95* (mean CPM = 263 ± 65 SD). About half of *Polistes* odorant receptors belong to an expanded clade of 9-exon subfamily ORs (Legan et al. 2021). In *P. fuscatus*, 9-exon ORs exhibited slightly lower mean transcription relative to the rest of the OR subfamilies (9-exon ORs mean CPM = 15.43; median CPM = 11.56; other ORs mean CPM = 22.98; median = 15.88; Wilcoxon rank sum test: *P* = 0.019; W = 4942). However, some 9-exon ORs at the base of the expanded subfamily clade were highly transcribed in both sexes (Fig. 2). L subfamily ORs were transcribed at higher levels than ORs in other subfamilies (L ORs mean CPM = 32.38; median CPM = 25.5; other ORs mean CPM = 17.59; median = 12.46; Wilcoxon rank sum test: *P* = 8.3e-05; W = 3511).

**Figure 2.**
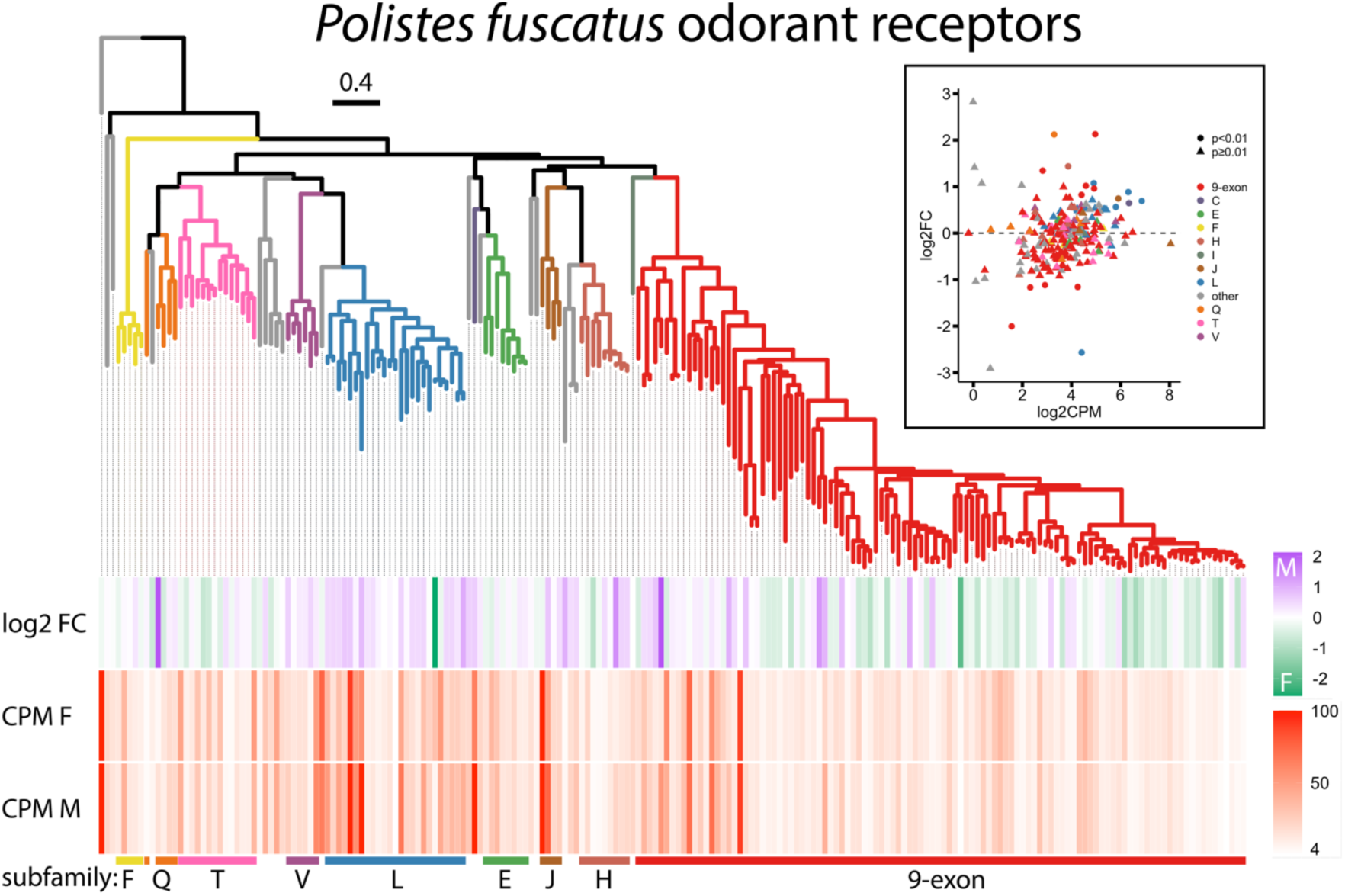
*P. fuscatus* odorant receptor gene tree, based on amino acid alignment, with gene transcription data for individual genes mapped to the tree’s branch tips. Scale bar at the top represents estimated divergence in amino acid substitutions per site. CPM represents the averages of counts per million per gene across six females (“CPM F”) and five males (“CPM M”). The upper right panel shows MA plot of *P. fuscatus* ORs. Tree branches and points in the MA plot (upper right panel) are color coded by odorant receptor subfamily.

Nineteen ORs were differentially transcribed between males and gynes at *P-adj* < 0.01 (Fig. 3). The proportion of OR genes that were differentially transcribed was not significantly different from the overall proportion of differentially transcribed genes (two-sided Fisher’s exact test; *P* = 0.1416; odds ratio = 0.695). Four ORs were significantly gyne biased: three 9-exon ORs (*PfusOr128*, *PfusOr143*, *PfusOr155*), and one OR in subfamily L (*PfusOr185*). Fourteen ORs were significantly male biased: five ORs in the L subfamily (*PfusOr179*, *PfusOr180*, *PfusOr186*, *PfusOr192*, *PfusOr195*), five ORs in the 9-exon subfamily (*PfusOr19*, *PfusOR94*, *PfusOr107*, *PfusOr221*, *PfusOr222*), and one OR each in subfamilies C (*PfusOr204*), H (*PfusOr48*), J (*PfusOr99*), and Q (*PfusOr98*).

**Figure 3.**
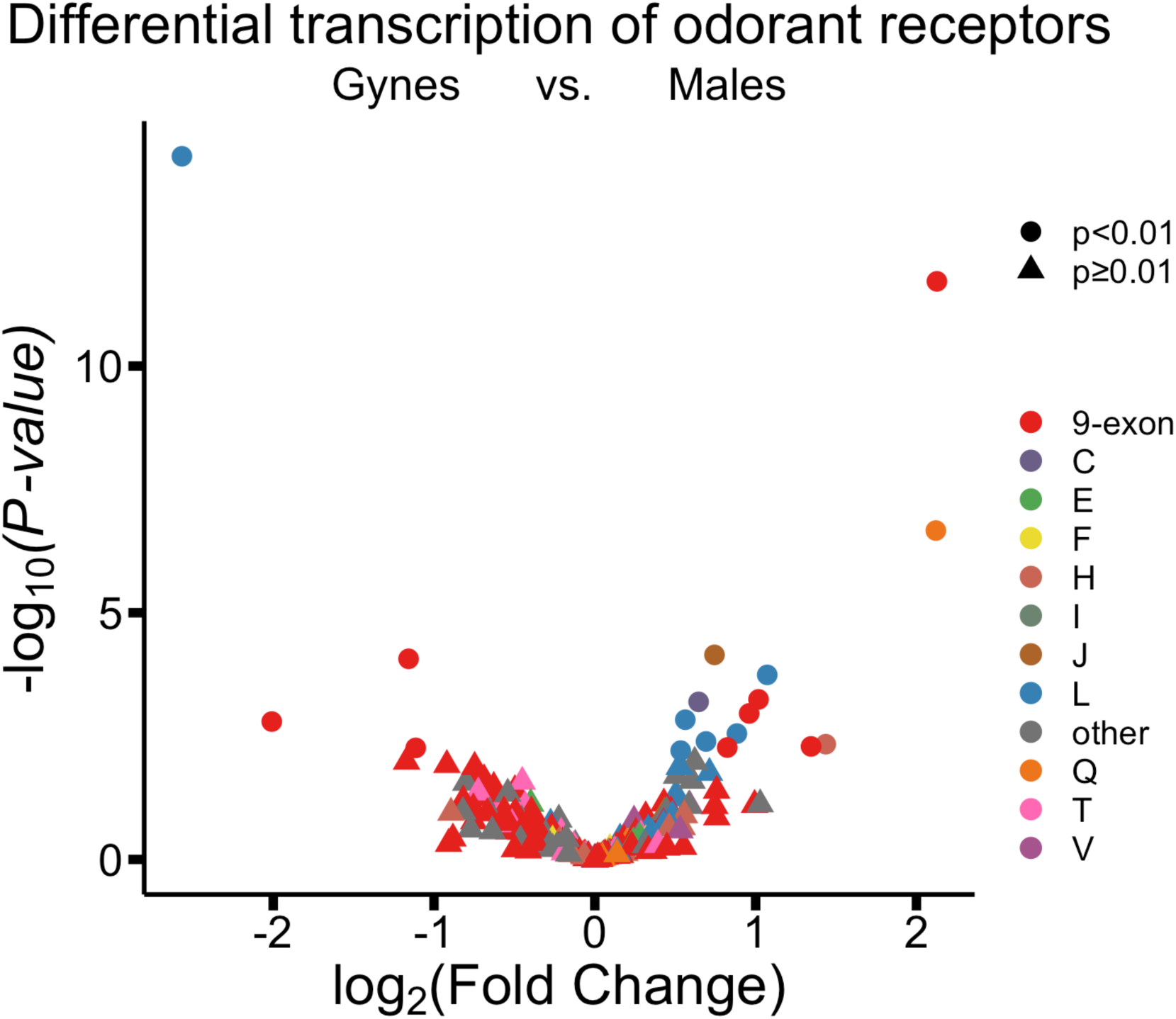
Volcano plot showing sexually dimorphic antennal gene transcription of the *P. fuscatus* OR genes, color-coded by OR subfamily. ORs with positive fold change values were biased towards male antennae, and those with negative fold change values were biased towards gyne antennae. OR genes represented by circles were differentially transcribed at *P-adj* < 0.01.

### Sexually dimorphic OR gene transcription among social Hymenoptera

In a honey bee, *Apis florea*, and three ant species, *Camponotus floridanus*, *Harpegnathos saltator*, and *Oocereae biroi*, antennal transcription of 9-exon ORs was found to be highly biased towards females (Zhou et al. 2012, 2015; Karpe et al. 2016; Fig. 4A). However, 9-exon OR transcription was not especially female biased in *P. fuscatus* compared to these species. The proportion of genes in the *P. fuscatus* 9-exon subfamily that showed any male bias (45/108) was substantially greater than in the other four social hymenopterans examined (two-sided Fisher’s exact test; *P* = 2.2e-16; odds ratio = 15.8; Fig. 4A). Values of sexually dimorphic transcription of 9-exon ORs in *P. fuscatus* were significantly different from those observed in the four other social Hymenoptera species (*P. fuscatus* 9-exon ORs mean log_2_(fold change) = -0.12; median log_2_(fold change) = -0.13; other species 9-exon ORs mean log_2_(fold change) = -3.25; median log_2_(fold change) = -3.12; Wilcoxon rank sum test: *P* < 2.2e-16; W = 65129). Moreover, a larger proportion (20/25) of the *P. fuscatus* L subfamily was male biased than that of the other four species (63/235) (two-sided Fisher’s exact test; *P* = 2.637e-07; odds ratio = 10.8; Fig. 4B), and values of sexual dimorphism were significantly different between *P. fuscatus* and other social insect species (*P. fuscatus* L ORs mean log_2_(fold change) = 0.27; median log_2_(fold change) = 0.47; other species L ORs mean log_2_(fold change) = -0.02; median log_2_(fold change) = -0.11; Wilcoxon rank sum test: *P* = 0.002; W = 2967).

**Figure 4.**
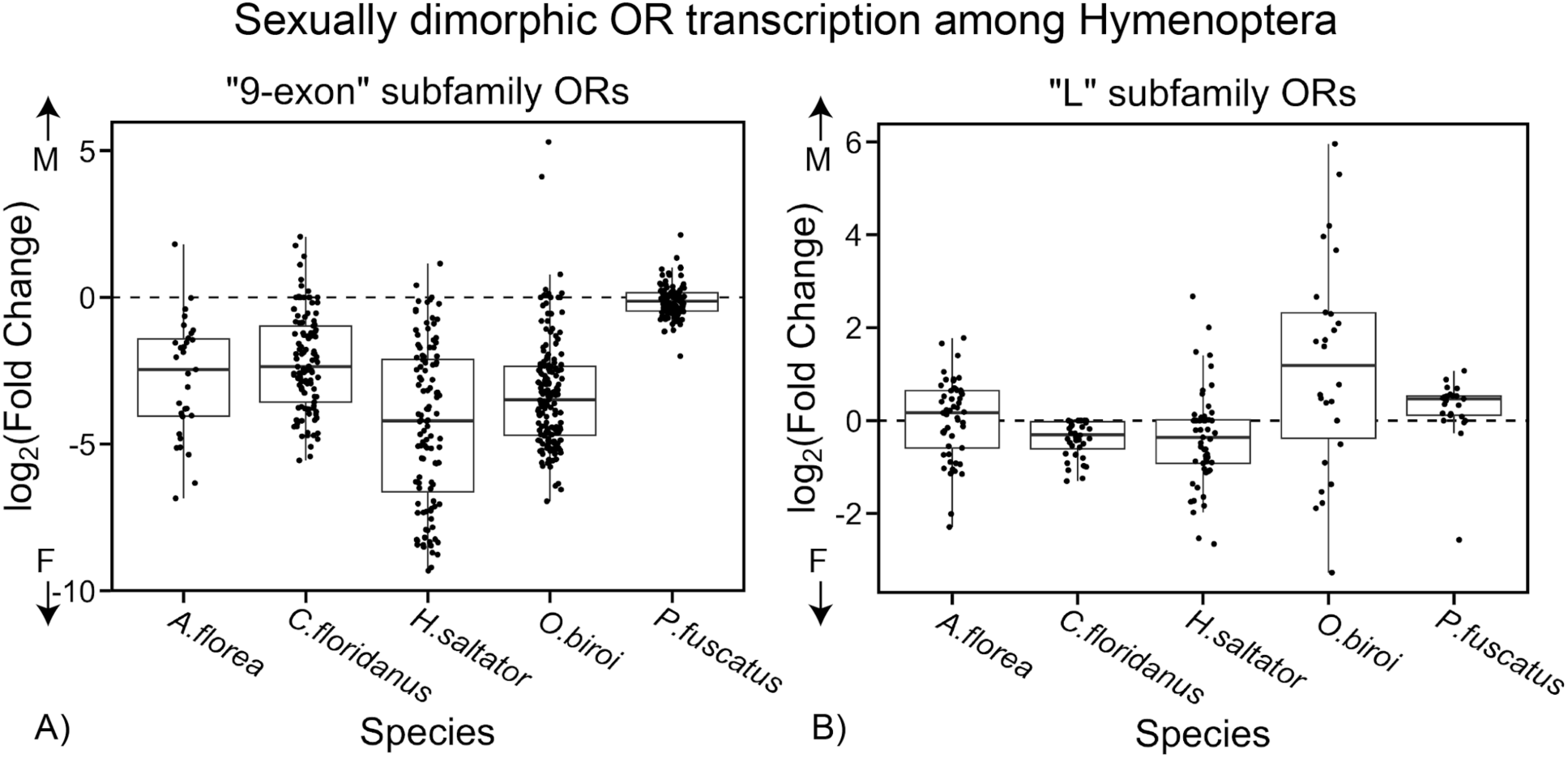
Sexually dimorphic antennal gene transcription as represented by log_2_(fold change) for (A) subfamily 9-exon and (B) subfamily L odorant receptors in the honey bee *Apis florea* (data from Karpe et al. 2016), three ant species (*Camponotus floridanus*, *Harpegnathos saltator*, and *Ooceraea biroi*; data from Zhou et al. 2015), and the paper wasp *Polistes fuscatus* (this study). ORs with positive fold change values were biased towards male antennae, and those with negative fold change values were biased towards female antennae. The median value is indicated by a solid black line within each box. Box lower and upper edges denote first and third quartiles. Whiskers extend to 1.5 times the inter-quartile range.

Finally, sexually dimorphic antennal OR gene transcription was mapped onto phylogenetic trees of the 9-exon and L OR subfamilies of a honey bee, two ant species, and a social wasp. In this phylogenetic context, *P. fuscatus* can be more clearly seen to depart from trends of female biased 9-exon OR gene transcription (Fig. 5A). Lineage-specific expansions of 9-exon ORs in *A. florea*, *C. floridanus*, and *H. saltator* tend to display worker biased antennal transcription (Fig. 5A). The relatively few male biased 9-exon ORs in *A. florea*, *C. floridanus*, and *H. saltator* are clustered in clades exhibiting relatively greater orthology near the base of the gene tree (Fig. 5A). Male biased ORs are scattered throughout the gene tree of L subfamily odorant receptors (Fig. 5B). *AfOr11*, ortholog of *Apis mellifera* queen pheromone receptor *AmOr11*, is within a basal clade of honey bee L subfamily ORs and does not exhibit clear orthology with ant or wasp ORs.

**Figure 5.**
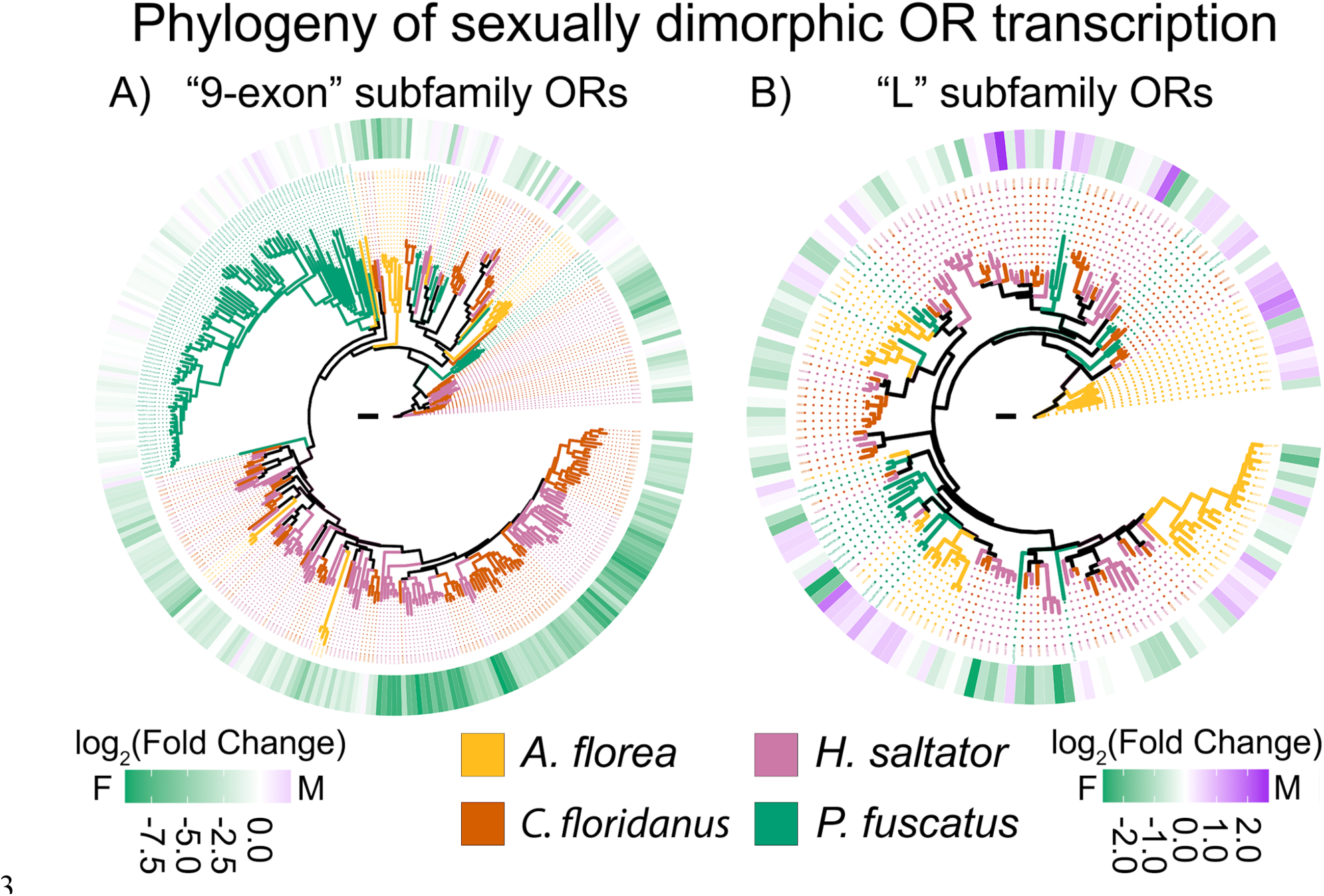
Phylogenetic reconstruction based on amino acid alignments of (A) 9-exon subfamily odorant receptors and (B) L subfamily odorant receptors from a honey bee *Apis florea* (data from Karpe et al. 2016), two ant species (*Camponotus floridanus*, *Harpegnathos saltator*; data from Zhou et al. 2012, 2015), and the paper wasp *Polistes fuscatus* (data from Legan et al. 2021, this study). Sexually dimorphic gene transcription data is mapped onto the trees as log_2_(fold change) where negative values (green) are female biased and positive values (purple) are male biased. Scale bars represent estimated divergence in amino acid substitutions per site (0.3).

## Discussion

### Sexual dimorphism in antennal transcription of genes related to metabolism and musculature

We identified patterns of differential gene transcription in the antennae of gyne and male social wasps, *Polistes fuscatus*. Bulk antennal mRNA sequencing revealed extreme sexual dimorphism in regulation of cytochrome P450 (CYP) genes, as well as genes putatively involved in muscular and glandular secretions, two morphological features that distinguish male and female antennae. Our finding of sexually dimorphic transcription of CYPs raises the intriguing possibility of sexually dimorphic odorant perception in paper wasps mediated by odorant degrading enzymes, though this will need to be empirically tested in future work. Odorant degrading enzymes help inactivate signals from lingering odorants and subsequently degrade the odorants in the sensillar lymph, safeguarding olfactory sensitivity by maintaining biologically relevant concentrations of odorants in the sensillar lymph (Ishida and Leal 2005, 2008; Durand et al. 2011; Leal 2013). Sexually dimorphic odorant degrading enzymes function in pheromone inactivation and degradation in the antennae of male silk moths and cotton leafworms (Vogt and Riddiford 1981; Durand et al. 2011). A honey bee worker biased CYP was posited to function in odorant degradation (Wanner et al. 2007b). CYPs also exhibited caste-specific transcription patterns in termites (Shigenobu et al. 2022). The frequent sexual dimorphism in CYP transcription observed in *P. fuscatus* antennae may reflect sexually dimorphic functions of CYPs in odorant inactivation and degradation. However, CYPs constitute a large and diverse gene family with multiple functions in degradation and synthesis, including toxin and xenobiotic degradation, modulation of juvenile hormone synthesis, cuticular hydrocarbon synthesis, and other functions (Sutherland et al. 1998; Danielson et al. 1998, 1999; Feyereisen 2006; Chung et al. 2009; Qiu et al. 2012).

Our analyses detected sexually dimorphic antennal transcription of several enzymes with putative functions in chemical synthesis, such as desaturation and elongation of fatty acids. Male biased fatty acid desaturase gene transcription is especially interesting because alkenes (unsaturated hydrocarbons) are among the cuticular hydrocarbons found to be male biased in other *Polistes* species (Beani et al. 2019). In multiple *Polistes* species, males tap females with their antennae during courtship, and they grip female antennae with their own during copulation (Mov. S1B). A unique attribute of the male *Polistes* antennae are their prehensile, curled antennae tips. Enrichment of GO terms related to muscle tissue may result from sexually dimorphic musculature of the antennae. Antennal secretory glands are sexually dimorphic in *Polistes dominula*, which led to the hypothesis that males secrete sex pheromones from their antennae (Romani et al. 2005). In other insects, for example in the cactophilic fly *Drosophila mojavensis*, males transfer chemical anti-aphrodisiacs to females which deter courtship by other males (Khallaf et al. 2020). Given the possibility of male-specific chemical signal synthesis combined with observations of male antennal gripping during courtship, chemical signal transfer during courtship by *Polistes* males should be investigated in the future.

### Sexual dimorphism in odorant receptor gene transcription

We identified 14 significantly male biased and four gyne biased odorant receptor genes. *Polistes* males and gynes likely rely on olfaction to locate mates. Male and female *P. exclamans* are attracted to hexane extracts of the opposite sex in a wind tunnel (Reed and Landolt 1990). *P. fuscatus* males are attracted to contents of the female venom gland of both *P. fuscatus* and *P. exclamans* and rely on cuticular chemicals of the female in mate compatibility recognition (Post and Jeanne 1984). Sexually dimorphic cuticular hydrocarbon profiles may be used in mate choice by *Polistes* males and females (Espelie and Hermann 1990; Layton et al. 1994; Beani et al. 2019; da Silva et al. 2021). Odorant receptors influencing mate choice may contribute to behavioral reproductive isolation, potentially influencing speciation (Brand et al. 2015, 2020; Xu and Shaw 2019). Both male and gyne *P. fuscatus* discriminate between conspecifics and heterospecifics of the closely related *P. metricus*, likely mediated in part by chemical cues or pheromones (Miller et al. 2019). Sexually dimorphic CHCs provide good candidate ligands for the sexually dimorphic OR genes identified in this study (Berson et al. 2019).

In social Hymenoptera, females are involved in more complex social interactions than males, while males usually have a more singular focus on mating. These differences in social behaviors appear to correlate with sexually dimorphic transcription of ORs in ants and bees. In ants, 9-exon subfamily ORs are enriched in antennae of female workers compared to males (Wanner et al. 2007b; Zhou et al. 2015; McKenzie et al. 2016). Indeed, in ants and social bees, the CHC-sensitive basiconic sensilla are completely absent on male antennae (Ågren and Hallberg 1996; Nakanishi et al. 2009; Nishino et al. 2009; Mysore et al. 2010; Ghaninia et al. 2018). On the other hand, CHC-sensitive antennal basiconic sensilla and associated neural architecture in the antennal lobe have been found in hornet males (Khodairy and Awad 2013; Couto et al. 2017, 2023). Our observation departs from previously reported patterns of OR transcription in ants and social bees. Most notably, the 9-exon subfamily that shows strongly female biased expression in honey bees and ants shows markedly less sexual dimorphism in transcription in *Polistes fuscatus*. While a narrow majority of *P. fuscatus* 9-exon ORs (63/108) trend toward female biased, many were male biased (Fig. 3). Moreover, the most male biased odorant receptor gene was a 9-exon receptor, *PfusOr94*, and in total five 9-exon receptors were significantly male biased compared to gynes. If 9-exon ORs detect social chemical signals in *Polistes*, as has been observed in ants, then how might *P. fuscatus* males utilize these receptors? *P. fuscatus* males, like males of other *Polistes* species, form aggregations away from the nest where they engage in weeks of social interactions (Mov. S1A). At these aggregations, male paper wasps might rely more on social chemical cues compared to males of other social insect species that do not form prolonged male aggregations.

We found that 9-exon OR genes were transcribed at marginally lower rates in both sexes compared to other subfamilies of ORs. The olfactory sensory neurons expressing 9-exon ORs likely innervate the T_B_ cluster of olfactory glomeruli, which tend to be individually small, though are collectively numerous (Couto et al. 2016, 2017; McKenzie et al. 2016). The smaller glomerular size suggests that each 9-exon receptor is likely expressed in a relatively smaller number of olfactory sensory neurons compared to receptors in other subfamilies. While a smaller number of neurons might express 9-exon ORs, tandemly arrayed ORs tend to be co-transcribed in clonal raider ant *O. biroi* olfactory sensory neurons (Brahma et al. 2023). Future work should investigate how this mechanism relates to bulk transcription patterns. Considering molecular evolutionary explanations for OR transcription, the 9-exon OR subfamily comprises several dynamically evolving tandem gene duplications in *Polistes* (Legan et al. 2021). Lower transcription levels are consistent with the hypothesis that most 9-exon OR genes originated from relatively recent gene duplication events, since younger genes tend to exhibit lower gene expression levels (Moutinho et al. 2022).

Our observation of generally male biased L subfamily OR transcription echoes results from social bees (Karpe et al. 2016). While solitary bees possess one-third the number of L subfamily receptors as honey bees, and the genome of the parasitoid wasp, *Nasonia vitripennis*, encodes few L subfamily ORs, the L subfamily is frequently expanded in social Hymenoptera (Zhou et al. 2015; Karpe et al. 2017; Legan et al. 2021). The honey bee L subfamily odorant receptor, *AmOr11*, is a drone biased sex pheromone receptor, responding to the queen substance component 9-ODA (Wanner et al. 2007b). The importance of this OR to males is reflected by the large size of the corresponding macroglomerulus MG2 in male antennal lobes (Sandoz 2006; Mariette et al. 2021). Several *H. saltator* L subfamily ORs responded to linear alkanes (Slone et al. 2017). While they tended to be male biased in mRNA abundance (20/25), L subfamily ORs were some of the highest transcribed ORs in both sexes. The most female biased odorant receptor gene was an L subfamily receptor, *PfusOr185*, indicating L subfamily ORs likely fulfill an important function for *P. fuscatus* males and females alike.

### Differences in gene transcription may underlie behavioral differences between life stages and castes

Sex-biased gene transcription of odorant receptors in social Hymenoptera has mainly been explored by comparing reproductive males to non-reproductive workers (Wanner et al. 2007b; Koch et al. 2013; Zhou et al. 2012; McKenzie et al. 2016). This comparison may represent differences in reproductive status rather than differences in sex per se. Recent transcriptomic analysis of the termite *Reticulitermes speratus* found that head gene transcription of reproductive males and reproductive females was more similar than between reproductive individuals and either males or females of soldier or worker classes (Shigenobu et al. 2022). A common tuning of the antennal olfactory system towards pheromones and chemical cues important for reproduction may reflect the relative lack of sexually dimorphic transcription of 9-exon ORs in *P. fuscatus*. As reproductive females move into different roles as foundresses and later queens, they may experience changes in OR transcript abundances. Differences in brain morphology and brain gene expression correlate with paper wasp caste differences and differences in social experience (O’Donnell et al. 2014; Toth et al. 2014; Jernigan et al. 2021; Uy et al. 2021). Comparisons of OR gene transcription between life stages will be useful to explore the relationship between wasp developmental stage and olfaction. Future work should also examine the gene regulatory mechanisms underlying sex- and caste-biased chemoreceptor gene transcription in *Polistes*, especially given that DNA methylation may not be an important factor shaping regulation of gene expression in social wasps (Patalano et al. 2015; Standage et al. 2016; Harrop et al. 2020; Miller et al. 2022).

### A role for social olfaction in male wasps

Due to alternate female dispersal strategies, where some females disperse long distances while others remain in the general vicinity of their nest of origin (Bluher et al. 2020), males and gynes may benefit by discriminating between related and unrelated mates. Fitness in both sexes benefits from outbreeding by reducing the rate of diploid male production, a costly phenomenon in haplodiploid social Hymenoptera (Boomsma et al. 2009). There is evidence both for (Ryan and Gamboa 1986) and against (Larch and Gamboa 1981; Post and Jeanne 1983b; de Souza et al. 2017a) the notion that *Polistes* males can recognize female nestmates, and this ability may vary between species. Ryan and Gamboa (1986) demonstrated that while *P. fuscatus* nestmate males and gynes were less aggressive towards each other than towards non-nestmates, they rarely mated. However, *P. versicolor* males did not avoid relatives or discriminate between female castes in laboratory mate choice assays (de Souza et al. 2017a; de Souza et al. 2021). Similarly, there is evidence in support of (Shellman-Reeve & Gamboa 1985) and against (Ryan et al. 1984) the claim that *P. fuscatus* males can recognize male nestmates. In *Polistes*, nestmate recognition phenotypes are complex cuticular hydrocarbon profiles (Gamboa et al. 1986; Espelie et al. 1994; Dani et al. 2001). Future work investigating how odorant receptors mediate nestmate recognition and other forms of olfactory recognition in *P. fuscatus* will be illuminating. The nature of social interactions within male mating aggregations is not well understood, and patterns of male biased transcription of putative social signal odorant receptors suggest a greater role for male chemosensory communication than previously recognized.

## Acknowledgments

This work was supported by the National Science Foundation [Graduate Research Fellowship Program grant number DGE-1650441 to A.W.L., CAREER grant number DEB-1750394 to M.J.S.], and National Institutes of Health [grant number DP2-GM128202 to M.J.S.]. The authors thank Dr. Jay Falk for helpful comments on an earlier version of the manuscript.

## Data Accessibility Statement

Data for this study were deposited at NCBI under BioProject ID PRJNA956068. The associated BioSamples are SAMN34203586-SAMN34203597. Twelve paired-end RNAseq libraries have been submitted to the NCBI Sequence Read Archive (SRA) to Sequence Read Archive (SRA) in association with BioProject ID PRJNA956068 under accessions SRR24182144-SRR24182155.

