## Supplementary Material for "Lineage-specific patterns of sexually dimorphic antennal transcription in the paper wasp *Polistes fuscatus*"

##### Table of Contents


| **Item** | **Description** | **Page number** |
|---|---|---|
| Supplementary Table S1 | Gene ontology enrichment analysis | **2** |
| Supplementary Figure S1 | MDS plot of sexually dimorphic gene transcription | **3** |
| Supplementary Figure S2 | MDS plot of sexually dimorphic gene transcription after removal of aberrant male sample "M50" | **4** |
| Supplementary Figure S3 | MA plot with *Polistes fuscatus* odorant receptors overlaid | **5** |

### Supplementary Tables

**Table S1.** Seventeen gene ontology term categories enriched for sexually dimorphic antennal gene transcription in *P. fuscatus*.

| GO ID | Term | Annotated | Significant | Expected | Classic Fisher |
|---|---|---|---|---|---|
| 0060047 | heart contraction | 48 | 8 | 6.8 | 3.20E-05 |
| 0042063 | gliogenesis | 87 | 13 | 12.32 | 0.00031 |
| 0045214 | sarcomere organization | 61 | 19 | 8.64 | 0.0005 |
| 0030239 | myofibril assembly | 89 | 30 | 12.6 | 0.00054 |
| 0003012 | muscle system process | 44 | 17 | 6.23 | 0.00208 |
| 0046434 | organophosphate catabolic process | 35 | 10 | 4.96 | 0.00253 |
| 0090175 | regulation of establishment of planar polarity | 23 | 8 | 3.26 | 0.0034 |
| 0001676 | long-chain fatty acid metabolic process | 12 | 6 | 1.7 | 0.00341 |
| 0098661 | inorganic anion transmembrane transport | 23 | 8 | 3.26 | 0.00468 |
| 0046716 | muscle cell cellular homeostasis | 29 | 10 | 4.11 | 0.00479 |
| 0007386 | compartment pattern specification | 13 | 6 | 1.84 | 0.00558 |
| 0007523 | larval visceral muscle development | 10 | 5 | 1.42 | 0.00764 |
| 0042593 | glucose homeostasis | 45 | 13 | 6.37 | 0.00768 |
| 0120036 | plasma membrane bounded cell projection organization | 912 | 146 | 129.16 | 0.00776 |
| 2000737 | negative regulation of stem cell differentiation | 18 | 7 | 2.55 | 0.00853 |
| 0006026 | aminoglycan catabolic process | 20 | 6 | 2.83 | 0.00978 |
| 0046068 | cGMP metabolic process | 17 | 5 | 2.41 | 0.00981 |

#### Supplementary Figures

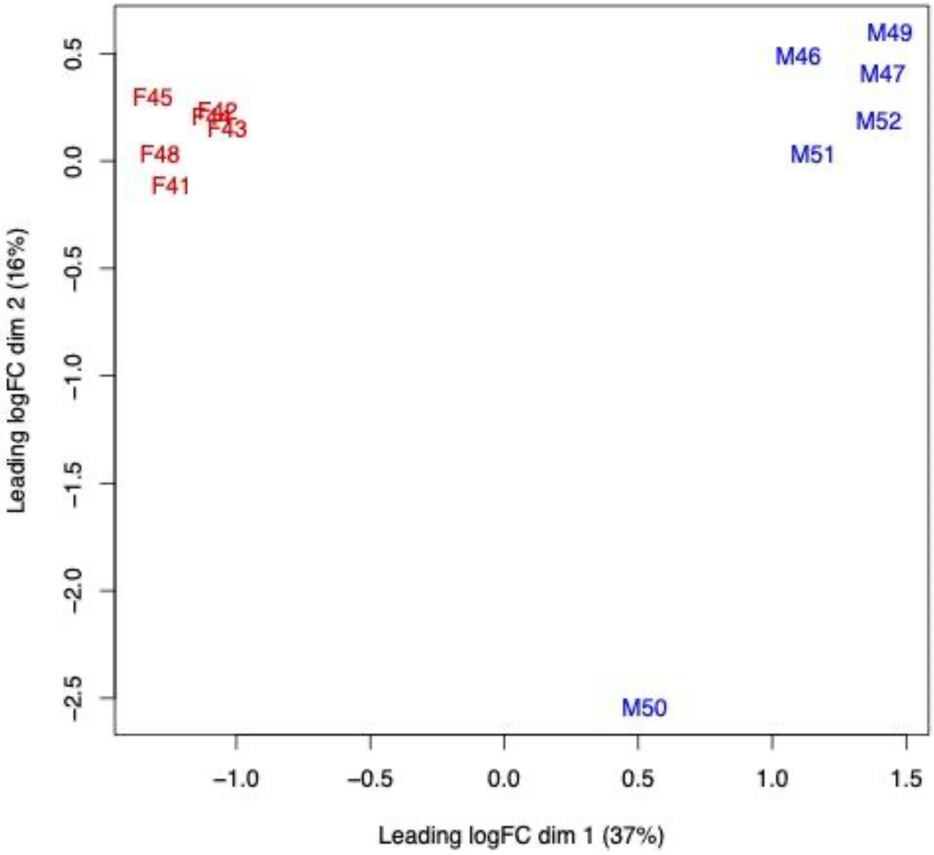


**Figure S1**. Multidimensional scaling plot of differences in antennal gene transcription between samples showing outlier sample “M50” which was removed from downstream analyses.

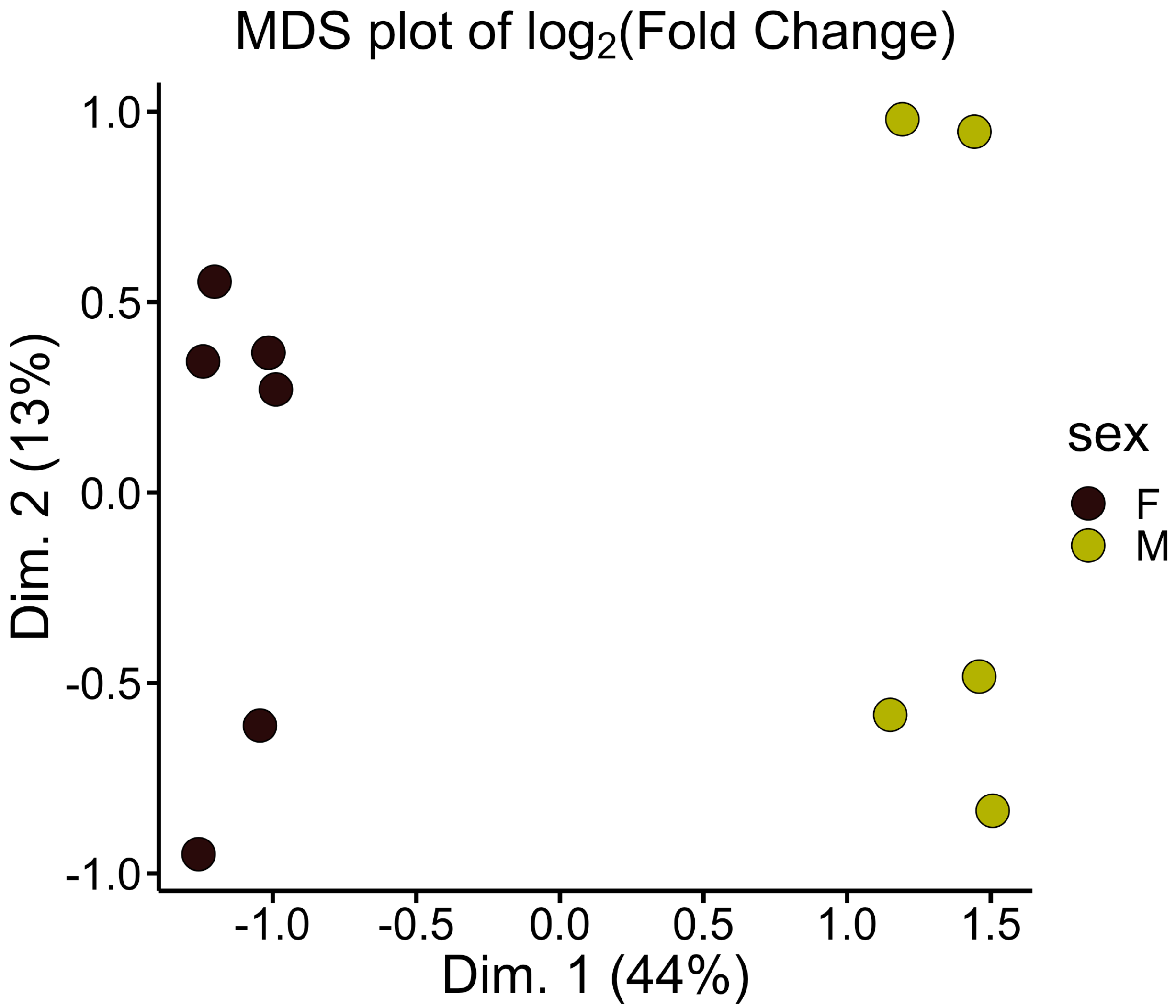


**Figure S2**. Multidimensional scaling plot of differences in antennal gene transcription between samples after removal of aberrant male sample “M50”. The overall $\log_2$(fold change) in mRNA read counts between the samples is roughly represented by the distances on the plot.

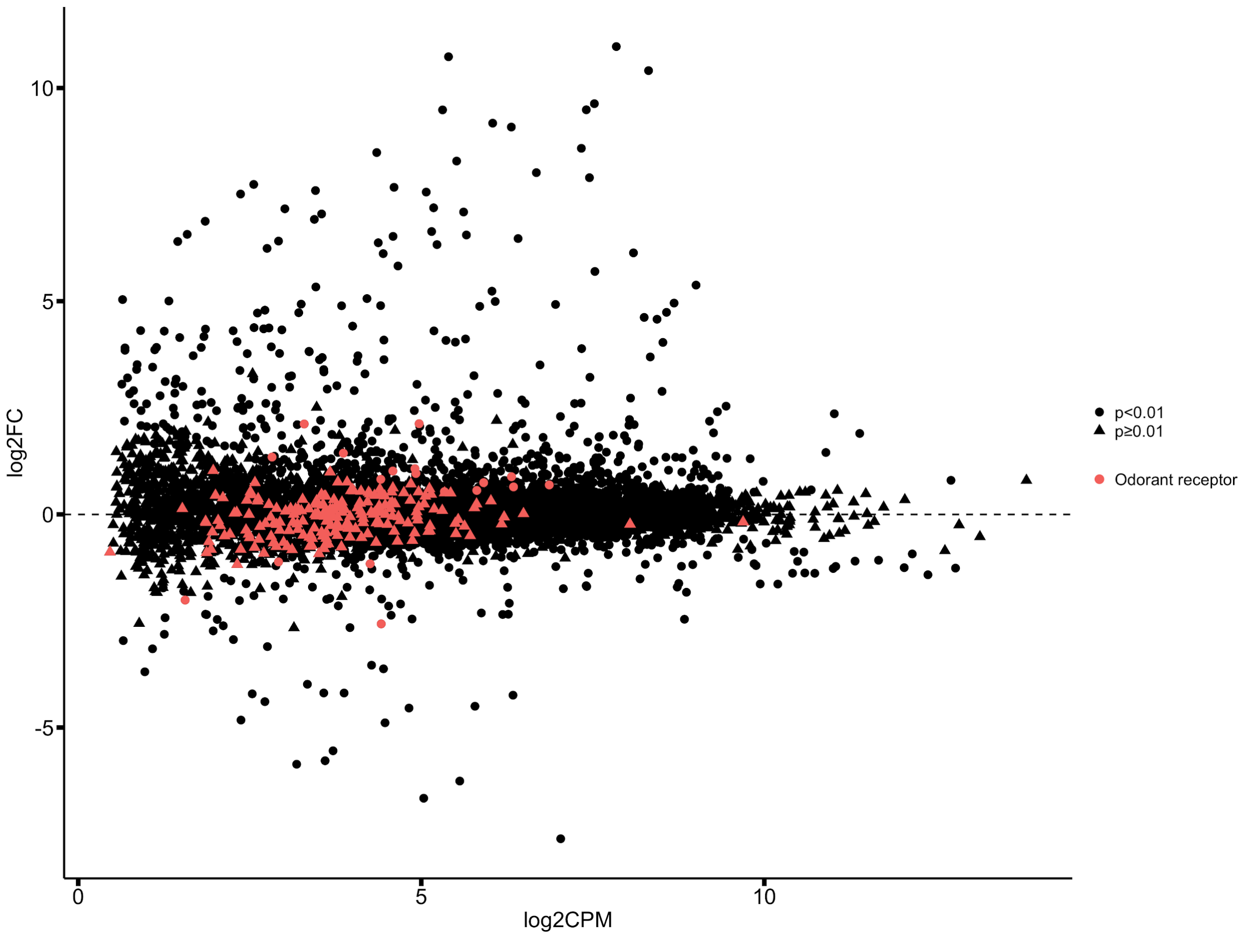


**Figure S3**. MA plot with *P. fuscatus* odorant receptors overlaid in red. The x-axis represents the base mean expression of each gene, measured as $\log_2$(counts per million). The y-axis represents fold change in gene transcription between males and gynes, measured as $\log_2$(fold change), where positive values indicate male biased gene expression and negative values indicate gyne biased gene expression.
